# An exploratory analysis of the current chemical regulations and guidelines from the perspective of endocrine disrupting chemicals using public resources

**DOI:** 10.1101/2020.10.08.331934

**Authors:** Bagavathy Shanmugam Karthikeyan, Janani Ravichandran, S. R. Aparna, Areejit Samal

## Abstract

The regulatory assessment of endocrine disrupting chemicals (EDCs) is complex due to the lack of a standardized definition of EDCs and validated testing criteria. In spite of these challenges, there is growing scientific interest in EDCs which has resulted in the rapid expansion of published literature on endocrine disruption upon chemical exposure. Here, we explore how academic research leading to curated knowledgebases can inform current chemical regulations on EDCs. To this end, we present an updated knowledgebase, DEDuCT 2.0, containing 792 potential EDCs with supporting evidence from 2218 research articles. Thereafter, we study the distribution of potential EDCs across several chemical lists that reflect guidelines for use or regulations. Further, to understand the scale of possible exposure to the potential EDCs present in chemical lists, we compare them with high production volume chemicals. Notably, we find many potential EDCs are in use across various product categories such as ‘Food additives and Food contact materials’ and ‘Cosmetics and household products’. Several of these EDCs are also produced or manufactured in high volume across the world. Lastly, we illustrate using an example how diverse information in curated knowledgebases such as DEDuCT 2.0 can be helpful in the risk assessment of EDCs. In sum, we highlight the need to bridge the gap between academic and regulatory aspects of chemical safety, as a step towards the better management of environment and health hazards such as EDCs.

## 1. Introduction

Endocrine disrupting chemicals (EDCs) are exogenous chemicals which interfere with the function of hormonal systems. They have been associated with hormone-related cancers and adverse effects in reproductive, developmental, and metabolic processes (Solecki et al., 2017; Swedenborg et al., 2009). They have also been observed to affect the immune system, the neurological system, and the liver (Solecki et al., 2017; Swedenborg et al., 2009; WHO/UNEP, 2013). EDCs are responsible for disease burden and impact on healthcare at an estimated annual cost of $340 billion in the USA and €163 billion in the European Union (EU) (The Lancet Diabetes & Endocrinology, 2019). Due to their hazard potential, EDCs and their adverse health effects on humans and wildlife have been investigated for more than three decades, and this information is documented in scientific literature including in published research articles, toxicological reports and regulatory guidelines (Darbre, 2019; WHO/UNEP, 2013). In fact, we will show here that the published evidence on EDCs in journal articles has seen significant growth over the last three decades (Figure 1a,b).

**Figure 1.**
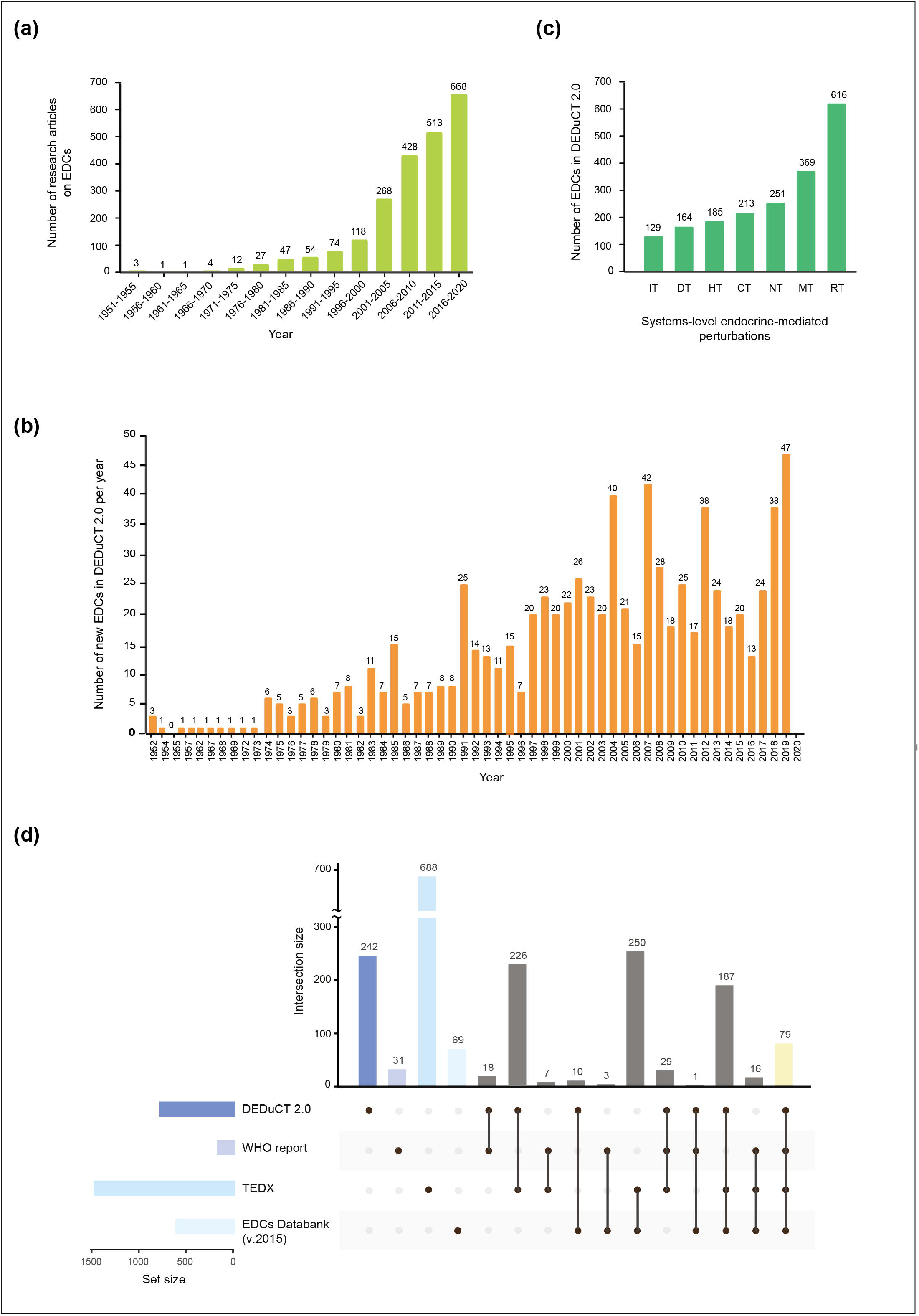
**(colour online): (a)** A chronological analysis of the corpus of 2218 published articles which form the supporting evidence for 792 potential EDCs in DEDuCT 2.0. **(b)** A plot of the number of new EDCs identified in published literature per year based on information compiled in DEDuCT 2.0. **(c)** Evidence for seven different systems-level perturbations from published experiments across 792 potential EDCs compiled in DEDuCT 2.0. **(d)** Comparison of the list of EDCs captured in DEDuCT 2.0 with three other resources. From the UpSetR plot, it is seen that 242 out of 792 potential EDCs in DEDuCT 2.0 are not captured in any other resource.

Despite this, several limitations and uncertainties challenge the risk assessment and regulation of EDCs (Darbre, 2019; Futran Fuhrman et al., 2015). Importantly, a standard (consensus) definition for EDCs can dictate the evidence needed for its identification among environmental chemicals (Zoeller et al., 2012; Futran Fuhrman et al., 2015; Solecki et al., 2017). In this direction, several definitions have been proposed and adopted by various regulatory agencies. However, clarity and standardization are yet to be achieved in EDCs research (Futran Fuhrman et al., 2015). This is also reflected in a recent comprehensive study commissioned by the European Parliament on endocrine disruptors and the current EU regulations on the subject (Demeneix et al., 2019). In particular, the report found gaps in the definition of EDCs, test requirements and guidelines for authorization of products in a number of categories such as cosmetics, drinking water and workers’ regulations (Demeneix et al., 2019). Another challenge to the regulation of EDCs is the wide range of factors to be considered in developing risk assessment criteria. In addition to defining the adverse effects, factors such as source and dosage of exposure need to be considered, all of which are aspects studied and documented in peer-reviewed articles in scientific journals. However, it is unknown to what extent this scientific literature is consulted during the development of risk assessment criteria and testing standards for EDCs. In fact, toxicity test guidelines have received criticism for having omitted several relevant endpoints which are captured in academic research (Ågerstrand et al., 2017).

The above-mentioned two observations, namely, the growth in the volume of scientific knowledge surrounding EDCs, and the perceived presence of gaps in the risk assessment and regulation of EDCs, have prompted the comparative analysis presented here. Specifically, we sought to understand how scientific knowledge from academic research can be useful in improving chemical regulation, with a focus on EDCs. We do so in three steps. We first aim to understand the increase in current knowledge surrounding EDCs via the increase in experimental evidence from published research articles. We then try to understand the presence of potential EDCs among chemicals we are exposed to in day-to-day life, for example, via the use of industrial and consumer products. Such an analysis of the presence of potential EDCs in man-made products is also a reflection of the regulations they are subject to. Lastly, we analyze which of these potential EDCs in human use are produced in large volume.

Previously, similar comparative studies (Neltner et al., 2013; Geueke et al., 2014) have been carried out for food, food additives and food contact substances, and these studies have highlighted the gaps in current regulation leading to presence of substances of concern in food and related products. However, previous studies were not specific to EDCs, and were also limited to a single category of substances. In this work, we have broadened the scope of our analysis and considered nine categories of substances based on the report commissioned by the European Parliament (Demeneix et al., 2019). On the other hand, we narrow our focus on EDCs, which are widely studied today and considered an emerging concern.

Overall, our three step analysis highlights the gap in transfer of knowledge between scientific research in academia on EDCs and chemical regulation. Further, this work shows how existing scientific knowledgebases can be helpful in framing criteria and strategies for risk assessment and regulation of EDCs. We believe the corpus of scientific information and evidence compiled in knowledgebases such as DEDuCT (Karthikeyan et al., 2019), will facilitate future regulation of EDCs and help deliver of safe consumer products to humankind.

## 2. Methods

### 2.1. DEDuCT 2.0 and other knowledgebases on EDCs

In order to explore the presence of EDCs in chemical lists used as guidelines or regulation, it is important to systematically compile potential EDCs along with their adverse effects. In this direction, we have previously created <u>D</u>atabase of <u>E</u>ndocrine <u>D</u>isrupting <u>C</u>hemicals and their <u>T</u>oxicity profiles (DEDuCT) which compiled information on 686 potential EDCs with supporting evidence from 1796 research articles published until February 2018 (Karthikeyan et al., 2019). In view of immense interest and growth of literature on the subject (Figure 1a,b), we here present an update, DEDuCT version 2.0, which compiles information on 792 potential EDCs with supporting evidence from 2218 research articles published until January 2020.

In Supplementary Text Section 1, we provide a detailed description of the four staged workflow designed to create DEDuCT by identifying potential EDCs with supporting evidence in published research articles (Supplementary Figures S1-S4; Supplementary Tables S1-S2). At the end of stage 1 of the workflow, we compiled 17134 potential research articles on EDCs published until January 2020 using a PubMed keyword search and information in three other resources on EDCs (Supplementary Figure S1). Of these 17134 research articles, 2837 articles were not evaluated in DEDuCT 1.0. Next, at the end of stage 2, we compiled 3996 research articles containing published experiments specific to humans or rodents on EDCs (Supplementary Figure S1). Of these 3996 research articles, 696 articles were not evaluated in DEDuCT 1.0. Thereafter, at the end of stage 3, we obtained 2047 chemicals which have been tested for endocrine disruption in humans or rodents in at least one of the 3996 research articles identified from stage 2 (Supplementary Figure S1). Of these 2047 chemicals, 421 chemicals were not evaluated in DEDuCT 1.0. Finally, at the end of stage 4, we compiled 792 potential EDCs with supporting evidence from 2218 research articles (Supplementary Figure S1). Notably, more than 100 potential EDCs in DEDuCT 2.0 were not captured in version 1.0 (Supplementary Table S2).

As mentioned in Supplementary Text Section 1, we have also captured significant pieces of information, as compiled in DEDuCT 1.0, such as observed endocrine-mediated endpoints and dosage information for EDCs in version 2.0. The entire compilation on 792 potential EDCs in DEDuCT 2.0 along with supporting evidence is publicly accessible via the web server: https://cb.imsc.res.in/deduct/.

We remark that DEDuCT compiles potential EDCs with supporting evidence specific to humans or rodents (Karthikeyan et al., 2019). In order to have a more comprehensive list of potential EDCs captured in published scientific literature, we considered three other prominent resources on EDCs, namely, the WHO report (WHO/UNEP, 2013), The Endocrine Disruption Exchange (TEDX; https://endocrinedisruption.org/), and the EDCs Databank version 2015 (Montes-Grajales and Olivero-Verbel, 2015) that contain information on 184, 1482, and 615 potential EDCs, respectively. The union of potential EDCs across the four resources, DEDuCT 2.0, WHO report, TEDX and EDCs Databank, was found to be 1856 chemicals (Figure 1d).

### 2.2. Compilation of chemical lists

To explore the extent to which current knowledge on EDCs in scientific literature is reflected in guidelines on chemical use or regulations worldwide, we systematically compiled such lists of chemicals that are part of inventories, guidelines and regulations from public resources. Through this exercise, we were able to compile 36 chemical lists which were broadly classified into two categories, namely, ‘Substances in use (SIU)’ and ‘Substances of concern (SOC)’ (Figure 2; Supplementary Text Section 2). In Supplementary Text Section 2, we provide a detailed description of these 36 chemical lists (L1-L36).

**Figure 2.**
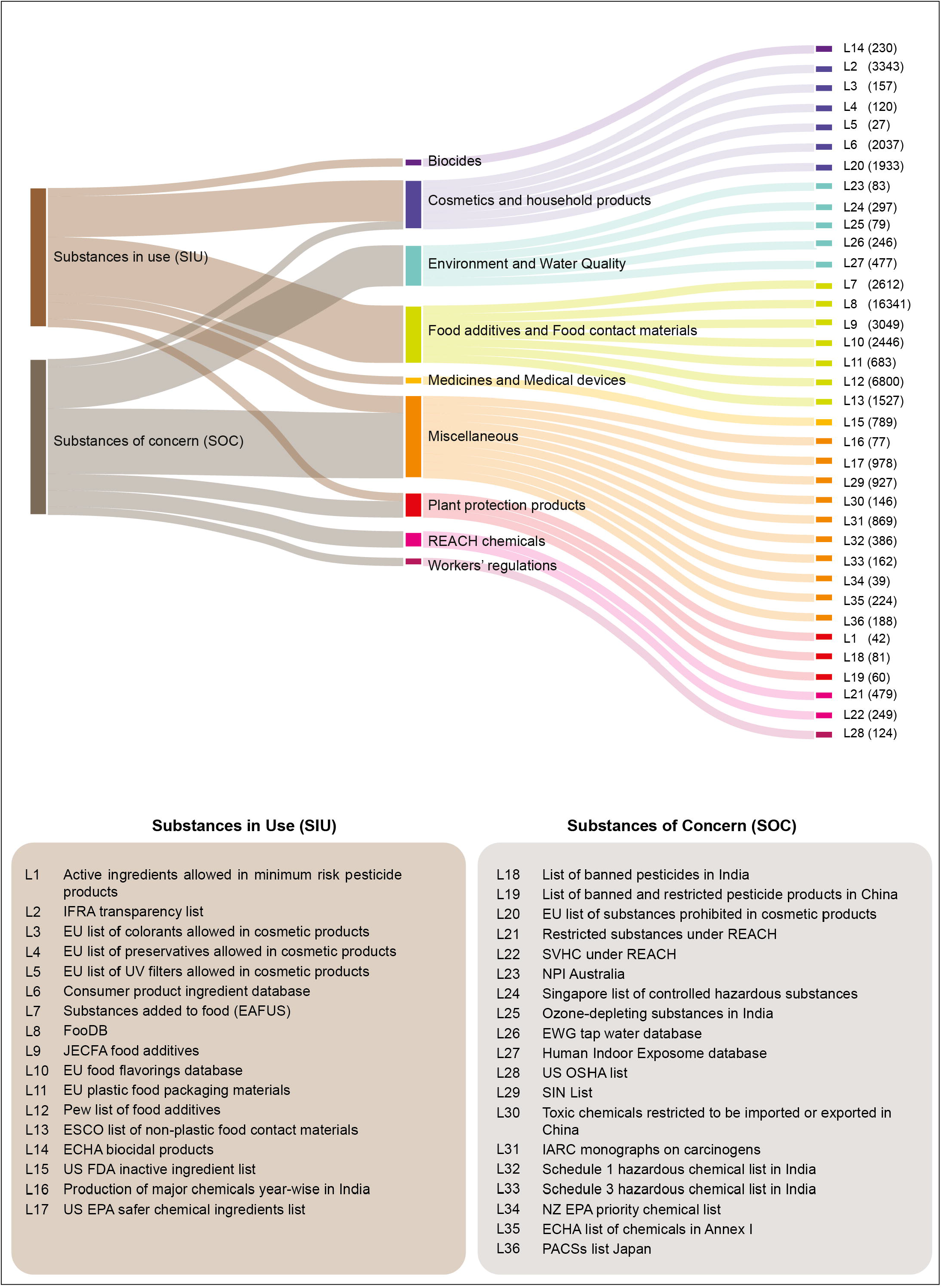
**(colour online):** Sankey plot showing the classification of 36 chemical lists that are part of inventories, guidelines and regulations obtained from public resources. The 36 chemical lists were broadly classified into two categories, namely, ‘Substances in use (SIU)’ and ‘Substances of concern (SOC)’. Based on chemical use or environmental source, the 36 chemical lists are further organized into 9 categories, namely, Plant protection products, Cosmetics and household products, Food additives and Food contact materials, Biocides, Medicines and Medical devices, REACH chemicals, Environment and Water Quality, Workers' regulations, and Miscellaneous. In this figure, the number of chemicals in each list is reported in parenthesis besides each list.

A list is considered a SIU list if it fulfills one of the following criteria: (a) It is an inventory of substances generally found to be in use in a certain product category; (b) It is a part of a guideline document, issued either by a government agency or an independent body, for safer product formulation; (c) It is a list of substances permitted for use in a certain product category, by a regulatory authority. Note that though inventories, regulations and guidelines, from where the 17 SIU lists were compiled, may have followed their own criteria to define the specific chemical lists, it is evident that the chemicals captured in these 17 SIU lists are in use in various consumer and industrial products.

An example of SIU list is the ‘L7 - Substances added to food (EAFUS)’ which is an inventory developed by the US Food and Drug Administration (FDA), and this list was previously known as Everything Added to Foods in the United States (EAFUS) (Figure 2; Supplementary Text Section 2). The L7 list contains 2612 unique chemicals which are used as food additives, color additives and other substances approved for specific use in food by the US FDA (Figure 2).

A list is considered a SOC list if it fulfills one of the following criteria: (a) It is an inventory of substances considered toxic, published either by a government agency or an independent body; (b) It is a list of substances monitored, restricted or banned for import, export or manufacture by a regulatory authority, due to their hazard potential.

An example of SOC list is the ‘L24 - Singapore list of controlled hazardous substances’ which is a chemical regulatory list compiled under the Schedule 2 of the Environmental Protection and Management Act of Singapore (Figure 2; Supplementary Text Section 2). The L24 list contains 297 hazardous substances (Figure 2).

Apart from the broad classification into SIU or SOC, we have also organized the 36 chemical lists into 9 categories based on the recent report commissioned by the European Parliament (Demeneix et al., 2019). These 9 categories include Plant protection products, Cosmetics and household products, Food additives and Food contact materials, Biocides, Medicines and Medical devices, REACH chemicals, Environment and Water Quality, Workers' regulations, and Miscellaneous (Figure 2). Note that we were able to find from public resources both SIU and SOC lists for only 3 out of these 9 categories (Figure 2).

For unequivocal analysis of chemicals in these 36 chemical lists representing guidelines or regulation, their respective Chemical Abstracts Service (CAS) identifiers (https://www.cas.org/) were used throughout this study.

### 2.3. Exploration of potential EDCs across chemical lists

Following the compilation of potential EDCs from four resources and 36 chemical lists, we have performed a three step systematic analysis to understand how potential EDCs are distributed across SIU and SOC lists.

First, we tried to identify any chemical overlap between the SIU and SOC lists. Upon finding a large chemical overlap between these two classes, we split the chemicals from the SIU and SOC lists into 3 groups (I-III). Group I consists of chemicals that are present only in 17 SIU lists, and not in any of the 19 SOC lists. Group II represents the list of chemicals that are present both in 17 SIU and 19 SOC lists. Group III represents the list of chemicals that are present only in 19 SOC lists, and not in any of the 17 SIU lists. We found 23483, 1139 and 3223 chemicals in group I, II and III, respectively (Figure 3a).

**Figure 3.**
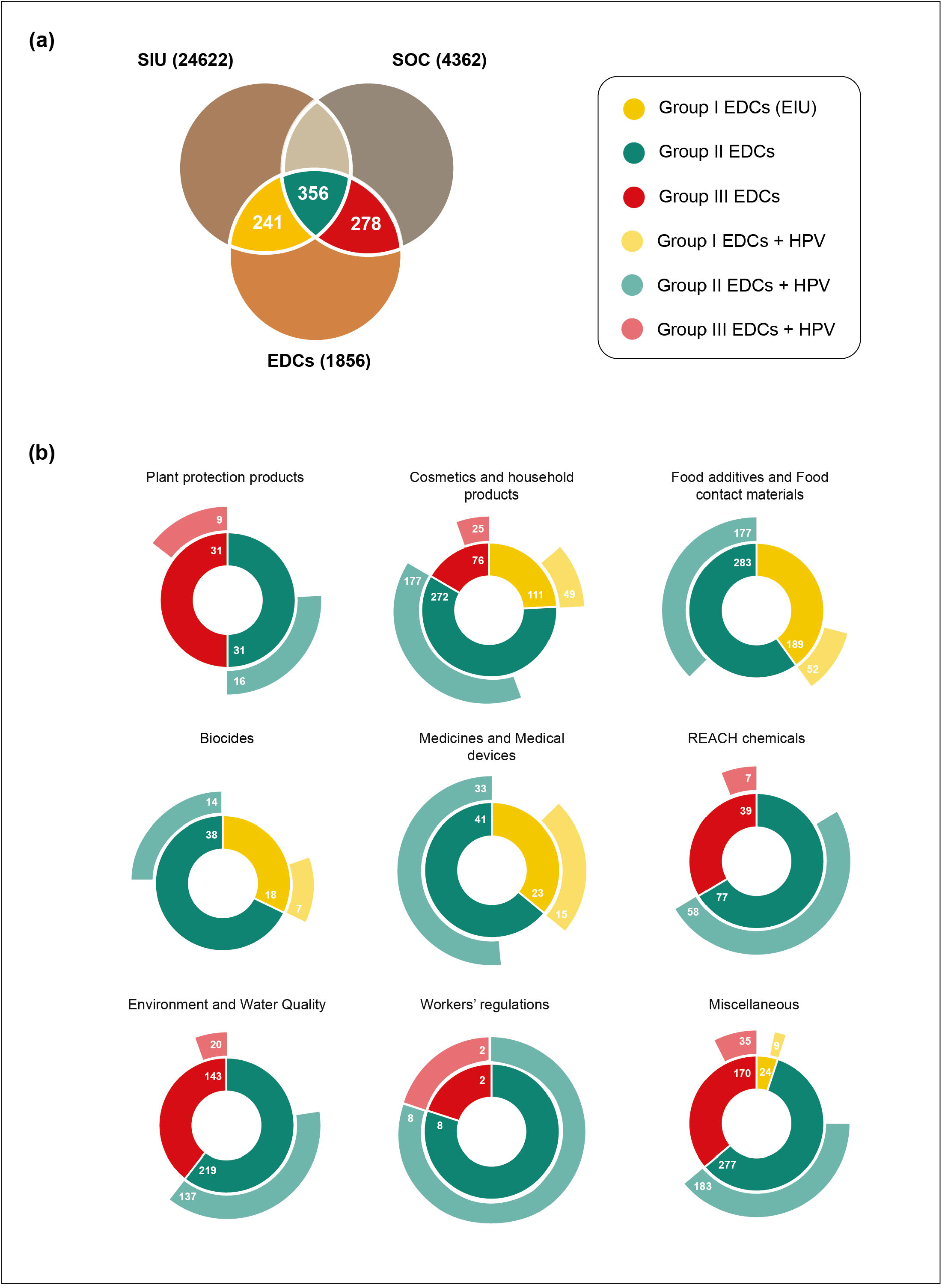
**(colour online):** Distribution of potential EDCs from four resources, namely, DEDuCT 2.0, WHO report, TEDX and EDCs Databank, across 36 chemical lists that are part of inventories, guidelines and regulations. **(a)** Venn diagram displaying the intersections of group I, II and III chemicals with potential EDCs (Methods). **(b)** Sunburst plot showing the distribution of potential EDCs across 9 categories of chemical lists. Within each category in this plot, the inner ring gives the number of potential EDCs in group I, II and III, and the outer ring gives the number of potential EDCs in group I, II and III that are also high production volume (HPV) chemicals.

Second, we compared the list of potential EDCs compiled from 4 resources, namely, DEDuCT 2.0, the WHO report, TEDX and EDCs Databank, with the group I chemicals. We refer to the list of potential EDCs in group I chemicals as group I EDCs or ‘EDCs in use (EIU)’ (Figure 3a). A similar comparison also led to group II EDCs and group III EDCs (Figure 3a).

Third, we compared the EIU list with the list of High Production Volume (HPV) chemicals to identify the potential EDCs in use which are produced or manufactured in high volume. For this analysis, we have compiled HPV chemicals from the union of two resources, namely, the United States High Production Volume (USHPV) database and the Organisation for Economic Co-operation and Development (OECD) High Production Volume (OECD HPV) list last updated on 2004. A similar comparison of the group II EDCs and group III EDCs was also performed with the HPV chemicals.

## 3. Results and discussion

### 3.1. DEDuCT 2.0 and growing research effort on EDCs

In previous work, we have built a unique knowledgebase, DEDuCT, containing information on 686 potential EDCs with supporting evidence from 1796 research articles (Karthikeyan et al., 2019). One goal of this work is to highlight the growing research effort in the academia on EDCs over the past decades. To create DEDuCT version 1.0 (Karthikeyan et al., 2019), we had mined and curated more than 16000 research articles published until February 2018 to finally obtain a corpus of 1796 articles containing supporting experimental evidence specific to humans or rodents for 686 potential EDCs. An analysis of this corpus of 1796 articles published until February 2018 found that the number of articles with supporting evidence on potential EDCs has significantly increased over the last three decades (Figure 1a). The continuous growth of literature on EDCs (Figure 1a) and community interest in DEDuCT 1.0 (Venkatasubramanian, 2019) served as motivation to perform a substantial update of our knowledgebase to include published scientific literature until January 2020.

For this work, we built an updated knowledgebase, DEDuCT version 2.0, with information on 792 potential EDCs with supporting experimental evidence from 2218 published research articles (Methods; Supplementary Text Section 1; Supplementary Tables S1-S2). In order to achieve the updated version 2.0, we had to mine and curate additional 3396 research articles which were published until January 2020. Essentially, we followed the same four staged workflow (Supplementary Figure S1) used to create DEDuCT 1.0 to create the updated version as described in Methods and Supplementary Text Section 1. The compiled information on 792 potential EDCs and additional information including supporting literature, systems-level perturbations, observed endocrine-mediated endpoints and corresponding dosage information is accessible via DEDuCT 2.0 web server at: https://cb.imsc.res.in/deduct.

A chronological analysis of the corpus of 2218 published articles which form the supporting evidence for 792 potential EDCs in DEDuCT 2.0 finds that there are 1181 articles published in the period 2011-2020, followed by 696 articles in the period 2001-2010, followed by 192 articles in the period 1991-2000 (Figure 1a). We remark that the corpus of 2218 research articles in DEDuCT 2.0 is likely to be a lower estimate of the accumulated scientific knowledge to date on EDCs; nevertheless, it is evident from Figure 1a that there has been significant growth in research on EDCs in the past three decades.

In addition, we leverage the 792 potential EDCs along with the associated supporting literature of 2218 research articles, to study the identification of new EDCs in the past decades. In Figure 1b, we show the number of new EDCs reported in published literature over the last 70 years. For this analysis, we consider a potential EDC captured in DEDuCT 2.0 to be identified for the first time in a particular year, if the earliest supporting experimental evidence for that EDC is from a research article published in that year. From Figure 1b, it is seen that the number of new EDCs identified in the scientific literature has slowly but surely increased on average over the past decades. These observations also align with the growth in scientific literature on EDCs.

A unique feature of our resource, DEDuCT 2.0, on EDCs is the compilation of observed 609 unique endocrine-mediated endpoints and their classification into 7 systems-level perturbations from supporting literature (Karthikeyan et al., 2019). We have also studied the available evidence for any of the 7 different systems-level perturbations across the 792 potential EDCs in DEDuCT 2.0 (Figure 1c). Of the 792 potential EDCs in DEDuCT 2.0, 616 EDCs have evidence for reproductive endocrine-mediated perturbations, 369 EDCs for metabolic perturbations and 251 EDCs for neurological perturbations (Figure 1c). This reflects that reproductive effects followed by metabolic effects may have been the main focus of the scientific investigations on EDCs.

Since DEDuCT compiles potential EDCs with supporting evidence specific to humans or rodents (Karthikeyan et al., 2019), we also considered three other resources on EDCs, namely, the WHO report (WHO/UNEP, 2013), TEDX and the EDCs Databank (Montes-Grajales and Olivero-Verbel, 2015) for the subsequent analysis (Methods). Figure 1d also gives an overview of unique and overlapping EDCs across the four resources. Specifically, 242 EDCs in DEDuCT 2.0 are not captured in any of the other three resources. In subsequent sections, we compare chemical lists pertaining to guidelines or regulations with the union of EDCs across the four resources which add up to 1856 potential EDCs (Figure 1d).

### 3.2. Distribution of potential EDCs across chemical lists

We have classified the chemicals in 17 SIU lists and 19 SOC lists into 3 groups I, II and III containing 23483, 1139 and 3223 chemicals, respectively (Methods). Thereafter, by comparing potential EDCs from four resources, namely, DEDuCT 2.0, WHO report, TEDX and EDCs Databank, we find 242, 356 and 278 potential EDCs in groups I, II and III, respectively (Figure 3a; Supplementary Table S3). Note that group II which is the intersection of chemicals present in SIU and SOC lists, contains more EDCs than groups I or III.

#### 3.2.1. Potential EDCs across substances in use

We designate the 242 potential EDCs among group I chemicals as EDCs in use (EIU) (Figure 3a; Supplementary Table S3). These 242 EIU are distributed across 5 of the 9 categories of chemical lists, and thus, pose a high risk of exposure (Figure 3a). Majority of EIU are found in 2 categories of chemical lists, namely, ‘Food additives and Food contact materials’ and ‘Cosmetics and household products’. Minority of EIU are found in 3 categories of chemical lists, namely, ‘Biocides’, ‘Medicines and Medical devices’ and ‘Miscellaneous’ (Figure 3b; Supplementary Table S3). Of the 242 EIU, DEDuCT 2.0 captures 119 potential EDCs along with supporting experimental evidence (Supplementary Table S3). Lastly, 6 EIU, namely, 2,4,5,2’,4’,5’-Hexabromobiphenyl, Coumestrol, Daidzein, Genistein, Pendimethalin and Zearalenone are captured in all four resources on EDCs (Supplementary Table S3).

#### 3.2.2. EDCs in use and high production volume chemicals

EIU produced in high volume can pose significant risk as humans are readily exposed to them through use of commercial products. Figure 3b gives the distribution of 63 EIU produced in high volume across 5 different categories of chemical lists (Methods; Supplementary Table S3). While none of EIU produced in high volume are captured in all four resources on EDCs, 7 EIU produced in high volume, namely, 4,4’-Dihydroxybiphenyl, 4-Hydroxybenzoic acid, 4-sec-Butylphenol, Chlorocresol, Monosodium glutamate, N,N’-Diphenyl-4-phenylenediamine and Sodium fluoride, are captured in three of the four resources on EDCs. These 7 EIU produced in high volume are found in 4 categories of chemical lists, namely, ‘Biocides’, ‘Cosmetics and household products’, ‘Food additives and Food contact materials’ and ‘Medicines and Medical devices’ (Figure 3b; Supplementary Table S3). Finally, 31 of the 63 EIU produced in high volume are captured in DEDuCT 2.0 (Supplementary Table S3).

From this analysis, it is evident that several EDCs in commercial use are also produced in high volume. The risk of exposure and associated hazard potential warrant an evaluation of these EIU produced in high volume, and framing appropriate risk assessment criteria will help such efforts. Later in this section, we illustrate how our knowledgebase, DEDuCT 2.0, on EDCs can aid in risk assessment.

#### 3.2.3. Potential EDCs across group II and III chemicals

There are 356 group II EDCs (Figure 3a) of which 211 are also HPV chemicals. Among the 356 group II EDCs, 46 are captured in all four resources on EDCs (Supplementary Table S3). Of these 46 group II EDCs, 28 are also produced in high volume. These 28 group II EDCs produced in high volume are distributed across 6 categories of chemical lists, namely, ‘Plant protection products’, ‘Cosmetics and household products’, ‘Food additives and Food contact materials’, ‘Environment and Water Quality’, ‘REACH chemicals’ and ‘Miscellaneous’ (Supplementary Table S3). Given the volume of production and their possible presence in commercial products, the risk of human exposure to these potential EDCs is a concern.

We next analyzed group III chemicals which are only present in SOC lists and found 278 potential EDCs among them (Figure 3a). Of these 278 group III EDCs, 5 chemicals, namely, Simazine, Linuron, Acetochlor, Vinclozolin, and Prochloraz, were found to be produced in high volume and captured in all four resources on EDCs (Supplementary Table S3). These 5 group III EDCs are distributed across 4 categories of SOC lists, namely, ‘Plant protection products’, ‘Cosmetics and household products’, ‘Environment and Water Quality’, and ‘Miscellaneous’ (Supplementary Table S3). These 5 potential EDCs in SOC lists are better monitored as they are produced in high volume in spite of known concern.

We further analyzed the distribution of HPV chemicals within potential EDCs in group II or III across the 9 categories of chemical lists (Figure 3b). Interestingly, we find that 33 out of 41 group II EDCs within ‘Medicines and Medical devices’ category are produced in high volume. Also, all group II or III EDCs within ‘Workers’ regulations’ category are produced in high volume indicating the risk of occupational exposure. Note that we were able to obtain only a single SOC list with 124 chemicals in the category of ‘Workers’ regulations’ of which 10 are potential EDCs produced in high volume (Figure 3b), and this analysis reveals current gap in regulation of occupational exposure to hazardous chemicals. Moreover, 3 potential EDCs namely, formaldehyde, ethylene oxide and methyl bromide, in the SOC list in ‘Workers’ regulations’ category are captured in three out of four resources on EDCs. Also, formaldehyde and ethylene oxide are present in 8 SIU lists suggesting potential risk of exposure from use of common products. A thorough evaluation of potential EDCs in 36 chemical lists (L1-L36), and incorporation of diverse information captured in scientific literature can improve safety assessment and regulation of EDCs.

#### 3.2.4. A case study of DEDuCT 2.0 in risk assessment of EDCs

To better understand how diverse information in a curated knowledgebase such as DEDuCT 2.0 can aid in chemical regulation, we present a case study for a potential EDC. We focused on 28 group II EDCs produced in high volume and captured in all four resources on EDCs including DEDuCT 2.0 (See preceding section). Of these 28 group II EDCs, ‘Dibutyl phthalate (CAS: 84-74-2)’ is a potential EDC present in 6 SIU lists and 7 SOC lists which are distributed across 5 categories, namely, ‘Cosmetics and household products’, ‘Food additives and Food contact materials’, ‘REACH chemicals’, ‘Environment and Water Quality’, and ‘Miscellaneous’. We next discuss the utility of DEDuCT 2.0 in risk assessment of chemicals using Dibutyl phthalate as an example.

According to the United States National Academy of Sciences, risk assessment involves four steps, namely, Hazard identification, Dose-response assessment, Exposure assessment, and Risk characterization (Council, 1983). Among the four resources on EDCs, notably, DEDuCT has compiled the observed endocrine-mediated endpoints and the dosage at which endpoints are observed, from published experiments specific to humans and rodents (Karthikeyan et al., 2019), and this information can aid in risk assessment process. DEDuCT 2.0 compiles supporting evidence on endocrine disruption upon Dibutyl phthalate exposure from *in vivo* experiments in rodents and *in vitro* experiments in humans which were published in 35 research articles.

For the first step in risk assessment, we used DEDuCT 2.0 to identify health hazards posed by Dibutyl phthalate. For Dibutyl phthalate exposure, DEDuCT 2.0 has compiled 81 endocrine mediated endpoints spanning 7 systems-level perturbations, namely, reproductive, developmental, metabolic, immunological, neurological, hepatic, and endocrine-mediated cancer (Supplementary Text Section 1). For the second step in risk assessment, one can use the dosage information complied in DEDuCT 2.0 for 81 endpoints observed upon Dibutyl phthalate exposure. In particular, we have analyzed the dosage information for Dibutyl phthalate compiled in DEDuCT 2.0 specific to endpoints observed in *in vivo* rodent studies using dosage unit as mg/kg/day (Supplementary Table S4). In these published *in vivo* rodent studies on Dibutyl phthalate, the test concentration range for different endpoints is 0.01-1000 mg/kg/day across compiled studies in DEDuCT 2.0, the lowest dose at which an adverse effect is observed in any of these studies is 0.01 mg/kg/day, and the highest dose at which no adverse effects are observed in any of the studies is 125 mg/kg/day (Supplementary Table S4). We remark that the compiled dosage information for Dibutyl phthalate in DEDuCT 2.0 is compatible with previous reports suggesting possible non-monotonic dose response for this chemical (Beausoleil et al., 2016).

The third step of exposure assessment involves the identification of routes, frequency and duration of exposure at the population level. Though DEDuCT 2.0 compiles information on environmental sources of potential EDCs, it does not capture their duration and routes of exposure. A possible expansion of the knowledgebase to include biomonitoring and epidemiological information for EDCs from published literature will further aid in exposure assessment and risk characterization; however, such an update of DEDuCT 2.0 requires significant effort beyond our current scope.

## 4. Conclusions

The number of chemicals introduced into the market for commercial purposes continues to be high. Adequate risk assessment strategies are needed now, more than ever, to cope with the increasing demand for safe product formulations. In general, regulatory standards and criteria differ across countries and this lack of standardization applies to the regulation of EDCs as well (Darbre, 2015; Clahsen et al., 2019). The regulatory assessment of EDCs is complex as there are several challenges and limitations associated with these substances (Darbre, 2015; Futran Fuhrman et al., 2015). In recent years there has been a rapid increase in endocrine disruption studies and the accumulation of knowledge surrounding EDCs (Figure 1a,b). However, regulatory assessments fall short due to the limitations and uncertainties in the risk assessment of EDCs (Darbre, 2015; Futran Fuhrman et al., 2015; Mihaich et al., 2017). This may be due to the lack of knowledge transfer from academic research to the regulatory assessment of EDCs.

In this work, we explored how knowledge on EDCs captured through academic research will help in risk and regulatory assessment of EDCs. We performed this analysis using three steps. Firstly, we have analyzed the increase in research efforts and knowledge on EDCs in past decades, and have captured newly available information into our unique resource DEDuCT 2.0 (Figure 1). Thus, DEDuCT 2.0 now compiles 792 potential EDCs along with 609 unique endocrine-mediated endpoints, spanning 7 systems-level perturbations. Secondly, we analyzed the distributions of 1856 potential EDCs compiled in DEDuCT 2.0 or three other resources, namely, WHO report, TEDX and EDCs Databank, across 36 chemical lists which are part of inventories, guidelines and regulations. Notably, we found several potential EDCs are distributed across diverse chemical lists, and further, some of these chemical lists with potential EDCs are in day-to-day product categories such as ‘Food additives and Food contact materials’ and ‘Cosmetics and household products’. Moreover, we classified the chemicals in SIU and SOC lists into groups I, II and III containing 23483, 1139 and 3223 chemicals, respectively, of which 242, 356 and 278, respectively, are potential EDCs (Figure 3a). Lastly, analysis of 242 group I EDCs with HPV chemicals found 63 group I EDCs in use which are also produced in high volume.

The presence of potential EDCs in these chemical lists (L1-L36) is a concern as humans are exposed to these potential EDCs via the use of industrial and consumer products. There is a need to incorporate endocrine disruption as a standard criterion in chemical risk assessment. Despite scientific efforts to evaluate the risks that EDCs pose, there is a gap in the transfer of knowledge to the policy planning level (Ågerstrand et al., 2017). Focused systematic review of these lists by regulatory agencies and non-governmental chemical advocacy groups, coupled with better incorporation of research data compiled in academic resources may help improve and strengthen chemical regulations and guidelines, and consequently, improve the safety of our products as well. Based on the extent and variety of information necessary for building regulatory standards, the utility of the WHO report, TEDX, and EDCs Databank in regulatory assessment may be limited. These resources lack the systematic compilation of observed adverse effects specific to endocrine disruption from published literature. The compilation of endocrine-mediated adverse effects along with dosage information in DEDuCT 2.0 may prove valuable in the risk assessment and regulation of EDCs as demonstrated using a case study for Dibutyl phthalate here.

As continuous scientific research brings new discoveries and a deeper understanding into the effects of chemicals, it becomes important to regularly monitor the substances permitted for use under various regulations, and substances generally found in use in products, through the same lens of scientific risk assessment, in order to restrict emerging substances of concern at the earliest. Inventories and independent guidelines of hazardous or toxic substances also need to be evaluated and brought under effective regulation. Information with a scientific basis is necessary to standardize criteria for this evaluation and risk assessment, especially in the case of a complex chemical class such as the EDCs. To this end, experimental evidence of endocrine disruption for potential EDCs compiled in knowledgebases could help in the early identification of hazardous substances, so that regulatory bodies can then streamline the process for safety testing, and in turn improve chemical safety standards.

## CRediT author contribution statement

**Bagavathy Shanmugam Karthikeyan:** Conceptualization, Data curation, Methodology, Formal analysis, Software, Writing – original draft, Writing – review & editing. **Janani Ravichandran:** Conceptualization, Data curation, Methodology, Formal analysis, Software, Visualization, Writing – original draft, Writing – review & editing. **S. R. Aparna:** Conceptualization, Data curation, Writing – original draft, writing – review & editing. **Areejit Samal:** Conceptualization, Investigation, Supervision, Project administration, Funding acquisition, Writing – review & editing.

## Acknowledgements

AS acknowledges support from the Science and Engineering Research Board (SERB) India through a Ramanujan fellowship (SB/S2/RJN-006/2014), the Department of Atomic Energy (DAE) India, and the Max Planck Society Germany through a Max Planck Partner Group in Mathematical Biology. The funders have no role in study design, data collection, data analysis, manuscript preparation or decision to publish.

## Supplementary Figures

**Supplementary Figure S1:** Detailed workflow for the compilation of potential EDCs in DEDuCT version 2.0.

**Supplementary Figure S2:** Schematic figure depicting the classification of the 609 endocrine-mediated endpoints into 7 systems-level perturbations in DEDuCT version 2.0.

**Supplementary Figure S3:** Classification of the 792 potential EDCs into 7 broad categories and 48 sub-categories based on their source in the environment. In this figure, the number of EDCs in DEDuCT version 2.0 contained in each category or sub-category is reported within the parenthesis.

**Supplementary Figure S4:** Classification of the 792 EDCs in DEDuCT version 2.0 into chemical kingdoms and chemical super-classes using ClassyFire. Of the 792 EDCs, 746 are organic and 46 are inorganic compounds. The 746 organic EDCs can be further classified into 19 super-classes while the 46 inorganic EDCs fall into 3 super-classes. The number of EDCs in each super-class is reported within the parenthesis.

## Supplementary Tables

**Supplementary Table S1:** List of 17134 research articles obtained at the end of stage 1 of the workflow that contain keywords related to EDCs. This list was compiled from PubMed, EDCs Databank (version 2015), TEDX and the WHO report. In stage 2 of the workflow, the articles were then filtered based on study type and test organism resulting in 3996 research articles specific to humans or rodents (Third column). Further, we compiled all the tested chemicals associated with the resulting articles and mapped them to their structural identifiers. Each tested chemical along with the associated literature was investigated for the presence of experimental evidence on endocrine-mediated endpoints upon exposure during stage 4 of the workflow. At the end of stage 4, we obtained 792 potential EDCs with supporting evidence from 2218 research articles (Fourth column).

**Supplementary Table S2:** The final list of 792 potential EDCs classified into 4 categories (I-IV) based on the type of supporting evidence for endocrine disruption in published experiments specific to humans or rodents.

**Supplementary Table S3:** The table lists the potential EDCs compiled from four resources, namely, DEDuCT 2.0, WHO report, TEDX and EDCs Databank, which are present in 36 chemical lists that are part of inventories, guidelines and regulations. The table also provides information on potential EDCs in group I, II and III chemicals. Further, a comparison is also performed with high production volume (HPV) chemicals.

**Supplementary Table S4:** Compilation from DEDUCT 2.0 of the endocrine-mediated endpoints and corresponding dosage information upon exposure of potential EDC, Dibutyl phthalate (CAS: 84-74-2), as reported in *in vivo* rodent (IVR) experiments in published literature. As a step in the risk assessment, the test and effective concentration expressed in the unit, mg/kg/day, from these published IVR studies was used to gain insights on dose-response upon chemical exposure.

