## Supplementary Text and Supplementary Figures for "An exploratory analysis of the current chemical regulations and guidelines from the perspective of endocrine disrupting chemicals using public resources"

**for**

### 1. DEDuCT version 2.0

The Database of Endocrine Disrupting Chemicals and their Toxicity profiles (DEDuCT) is an online knowledgebase that compiles information on potential Endocrine Disrupting Chemicals (EDCs) (Karthikeyan et al., 2019). Notably, DEDuCT compiles supporting evidence for potential EDCs from published research articles that have reported endocrine disruption specific to humans or rodents upon chemical exposure in *in vivo* or *in vitro* experiments (Karthikeyan et al., 2019). DEDuCT version 1.0 released on 25 April 2019 compiled information on 686 potential EDCs with supporting evidence from 1796 research articles (Karthikeyan et al., 2019). In view of the steadily expanding literature on EDCs, we present here a significant update, DEDuCT version 2.0 released on 2 October 2020, which compiles information on 792 potential EDCs with supporting evidence from 2218 research articles. DEDuCT 2.0 is also accessible at: https://cb.imsc.res.in/deduct.

#### 1.1. Workflow for the compilation of EDCs

To compile the potential EDCs in DEDuCT, we have previously developed a four staged workflow (Supplementary Figure S1) to identify chemicals with supporting evidence of endocrine disruption in published literature specific to humans or rodents (Karthikeyan et al., 2019). The four stages in the workflow for the compilation of potential EDCs are summarized below.

##### 1.1.1. Literature mining

As the first step in stage 1 of the workflow, we mined PubMed abstracts (https://pubmed.ncbi.nlm.nih.gov/) using keywords related to EDCs. Note that we had mined PubMed abstracts until February 2018 for creating the earlier release DEDuCT 1.0. For the current release DEDuCT 2.0, this keyword search was last carried out on 13 January 2020, and has led to a compilation of 19723 research articles that are likely to contain information on EDCs. In the next step, we have compiled published research articles associated with EDCs listed in other publicly available resources, namely, the World Health Organization (WHO) report (WHO/UNEP, 2013), the Endocrine Disruption Exchange (TEDX version September 2018; https://endocrinedisruption.org/) and EDCs Databank version 2015 (Montes-Grajales and Olivero-Verbel, 2015), and this resulted in 337, 1166, and 456 published research articles, respectively. Finally, we obtained 17134 research articles that contain keywords related to EDCs through manual filtration of abstracts at the end of stage 1 (Supplementary Figure S1; Supplementary Table S1).

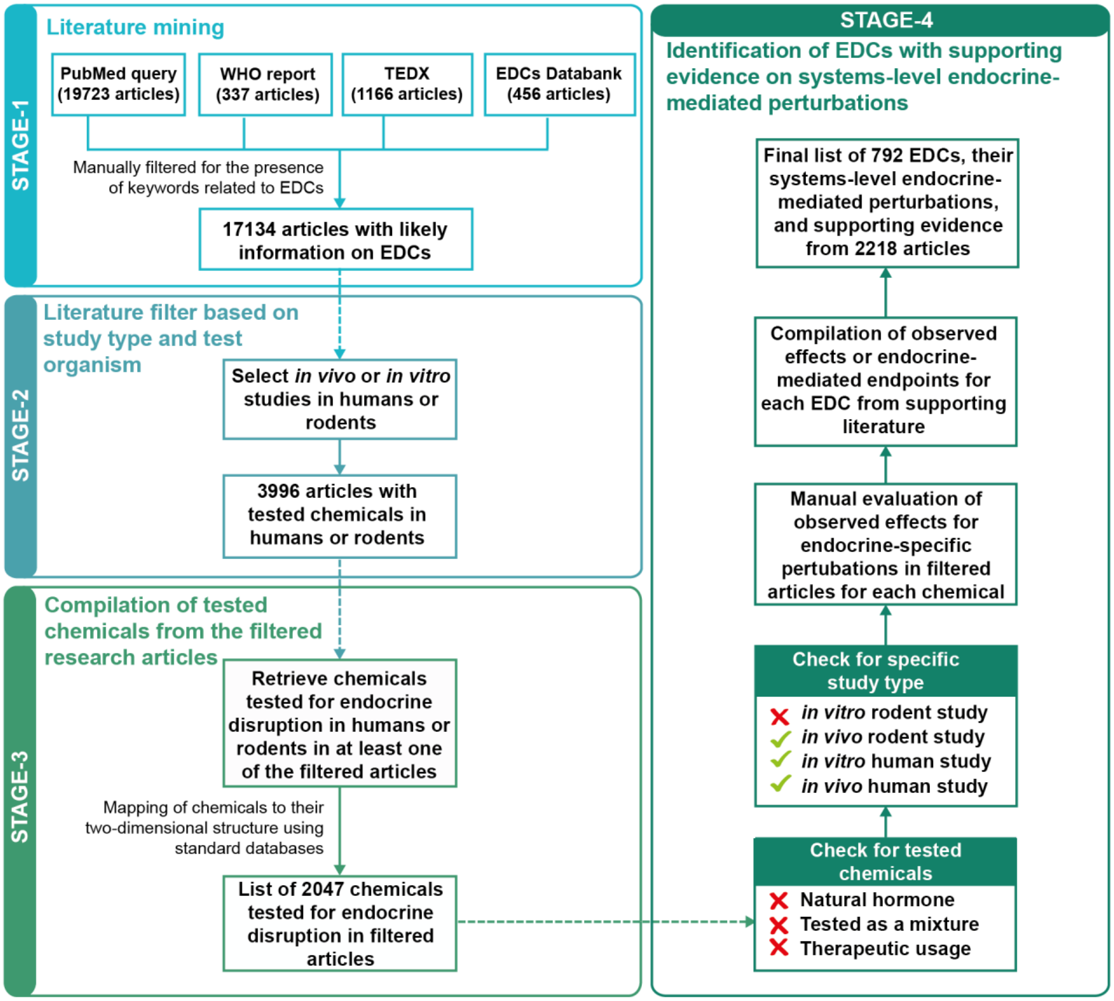


**Supplementary Figure S1:** Detailed workflow for the compilation of potential EDCs in DEDuCT version 2.0.

##### 1.1.2. Literature filter based on study type and test organism

In stage 2 of the workflow, we have screened 17134 research articles compiled in stage 1 to identify the studies based on *in vivo* and *in vitro* experiments in humans or rodents. We have removed any studies that infer potential endocrine disruption based on receptor-based binding assays or *in silico* methods. At the end of stage 2, we compiled 3996 research articles containing published experiments on EDCs specific to humans or rodents (Supplementary Figure S1; Supplementary Table S1).

##### 1.1.3. Compilation of tested chemicals from the filtered research articles

In stage 3 of the workflow, we collected all the tested chemicals associated with the resulting research articles from stage 2. Further, we mapped these chemicals to their two-dimensional (2D) structure identifiers using standard chemical databases. This resulted in 2047 chemicals with standard structure identifiers which have been tested for endocrine disruption in humans or rodents in at least one of the research articles identified at the end of stage 2 (Supplementary Figure S1).

##### 1.1.4. Identification of EDCs with supporting evidence on systems-level endocrine-mediated perturbations

In stage 4 of the workflow, we have assessed each tested chemical for the presence of significant observed effects in published experiments on endocrine disruption upon exposure (Supplementary Figure S1). As a first step, we have excluded the tested chemicals if, the chemicals are natural hormones, or the chemicals were tested as a part of a mixture, or the chemicals were tested for therapeutic usage in the published literature. Additionally, we have ignored the supporting evidence from published experiments performed on *in vitro* rodent test systems. Subsequently, we have evaluated the tested chemicals for the presence of significant observed effects or endpoints related to endocrine disruption such as changes in morphology, physiology, reproduction, growth and development, and lifespan. A chemical is identified as an EDC if at least one such endocrine-mediated endpoint is observed upon exposure in published experiments. At the end of stage 4, we identified 792 potential EDCs with strong supporting evidence for endocrine disruption from 2218 research articles specific to humans or rodents (Supplementary Table S1-S2).

Further, we have manually standardized the terms used to report observed effects upon endocrine disruption in published research articles, and this resulted in the compilation of 609 endocrine-mediated endpoints. These endocrine-mediated endpoints were further classified into 7 systems-level perturbations based on the major biological processes controlled by the human endocrine system, namely, Endocrine-mediated cancer (CT), Reproductive (RT), Developmental (DT), Metabolic (MT), Immunological (IT), Neurological (NT), Hepatic (HT) endocrine-mediated perturbations (Supplementary Figure S2). These unified terms can serve as standard biological vocabulary describing the toxicity profiles of EDCs. In addition to the compilation of endocrine-mediated endpoints, we have compiled the dosage range at which endocrine disruption was observed for individual EDCs in the published experiments in the supporting literature. Further, the compiled dosage units were uniformized, thereafter, providing the test and effective dosage for each EDC that could be used to determine dose-response measures.

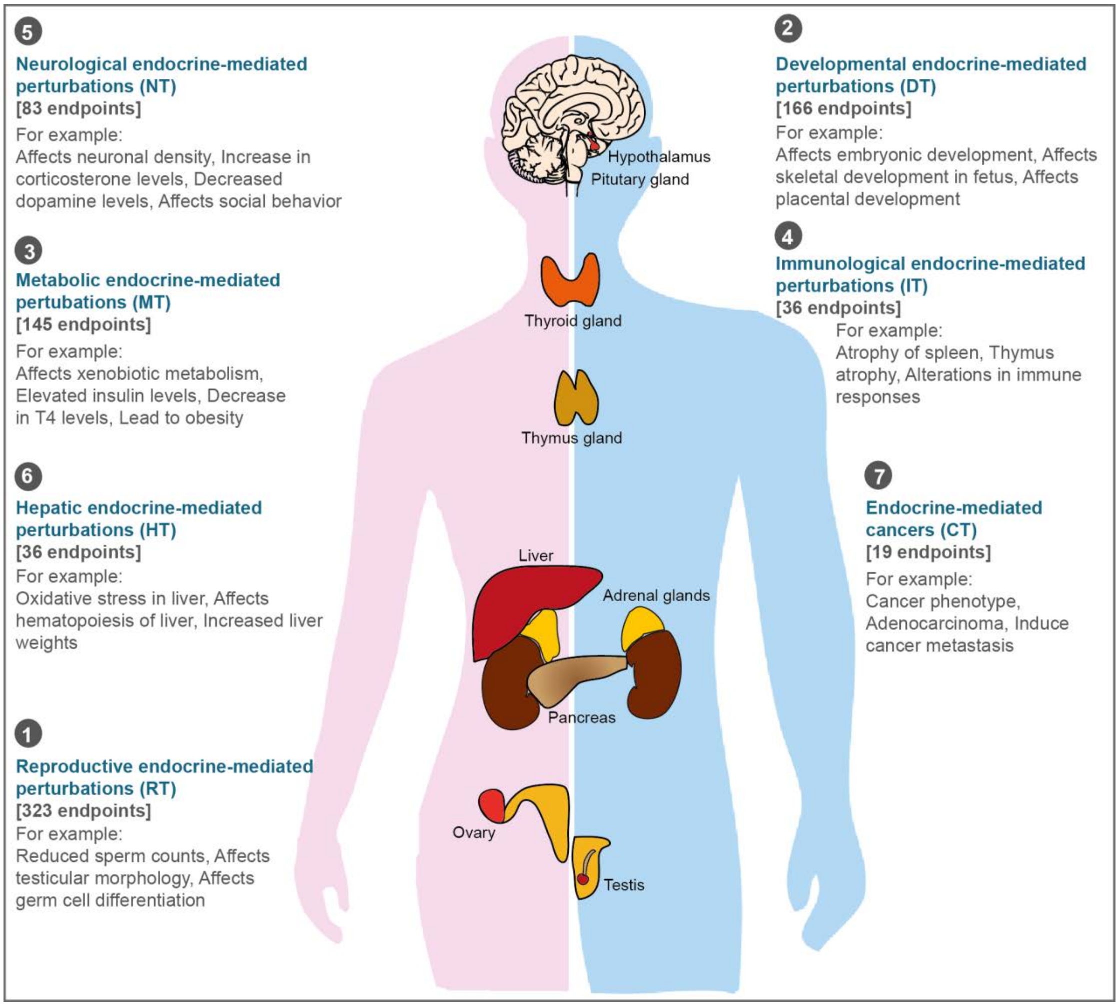


**Supplementary Figure S2:** Schematic figure depicting the classification of the 609 endocrine-mediated endpoints into 7 systems-level perturbations in DEDuCT version 2.0.

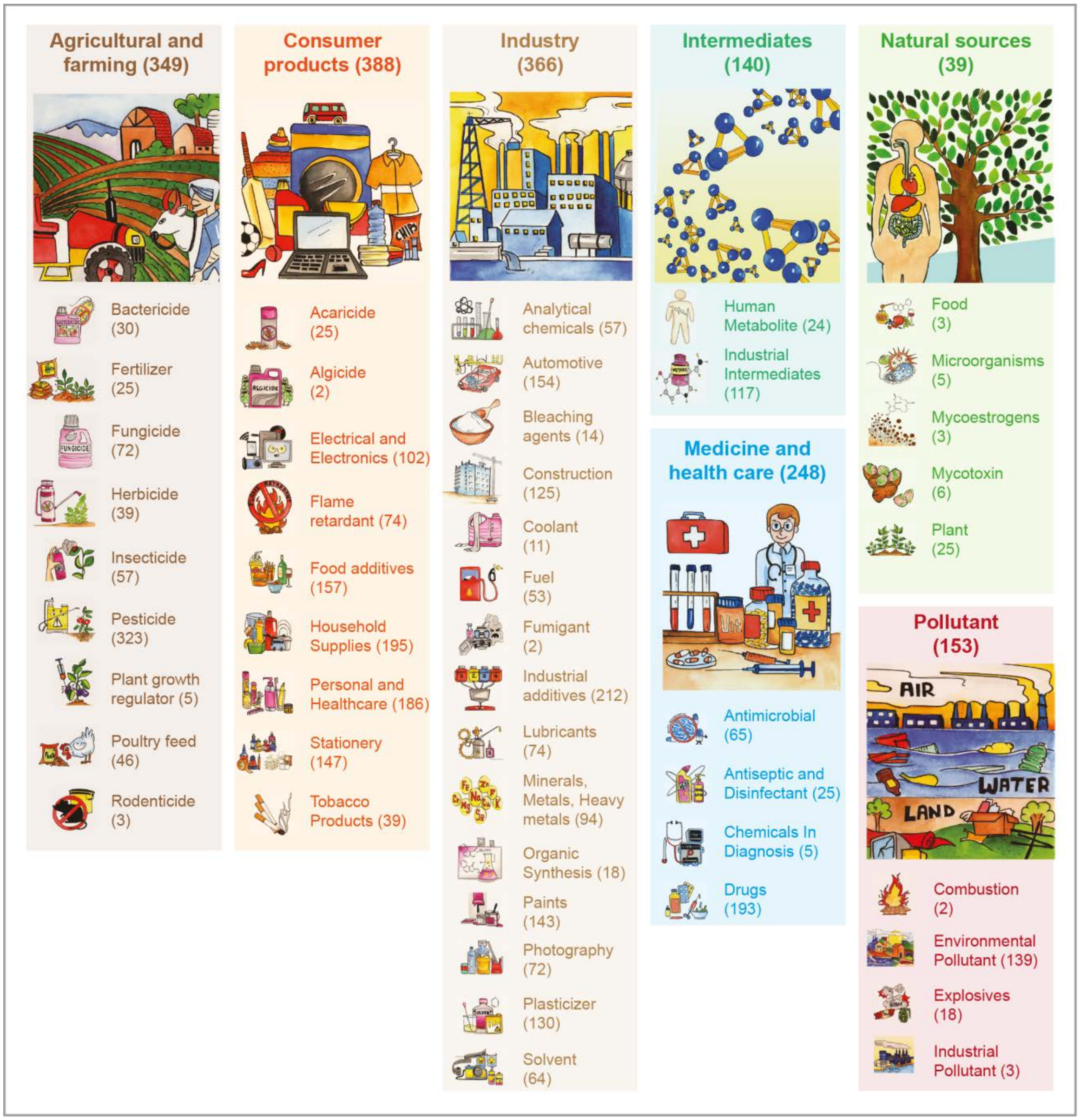


**Supplementary Figure S3:** Classification of the 792 potential EDCs into 7 broad categories and 48 sub-categories based on their source in the environment. In this figure, the number of EDCs in DEDuCT version 2.0 contained in each category or sub-category is reported within the parenthesis.

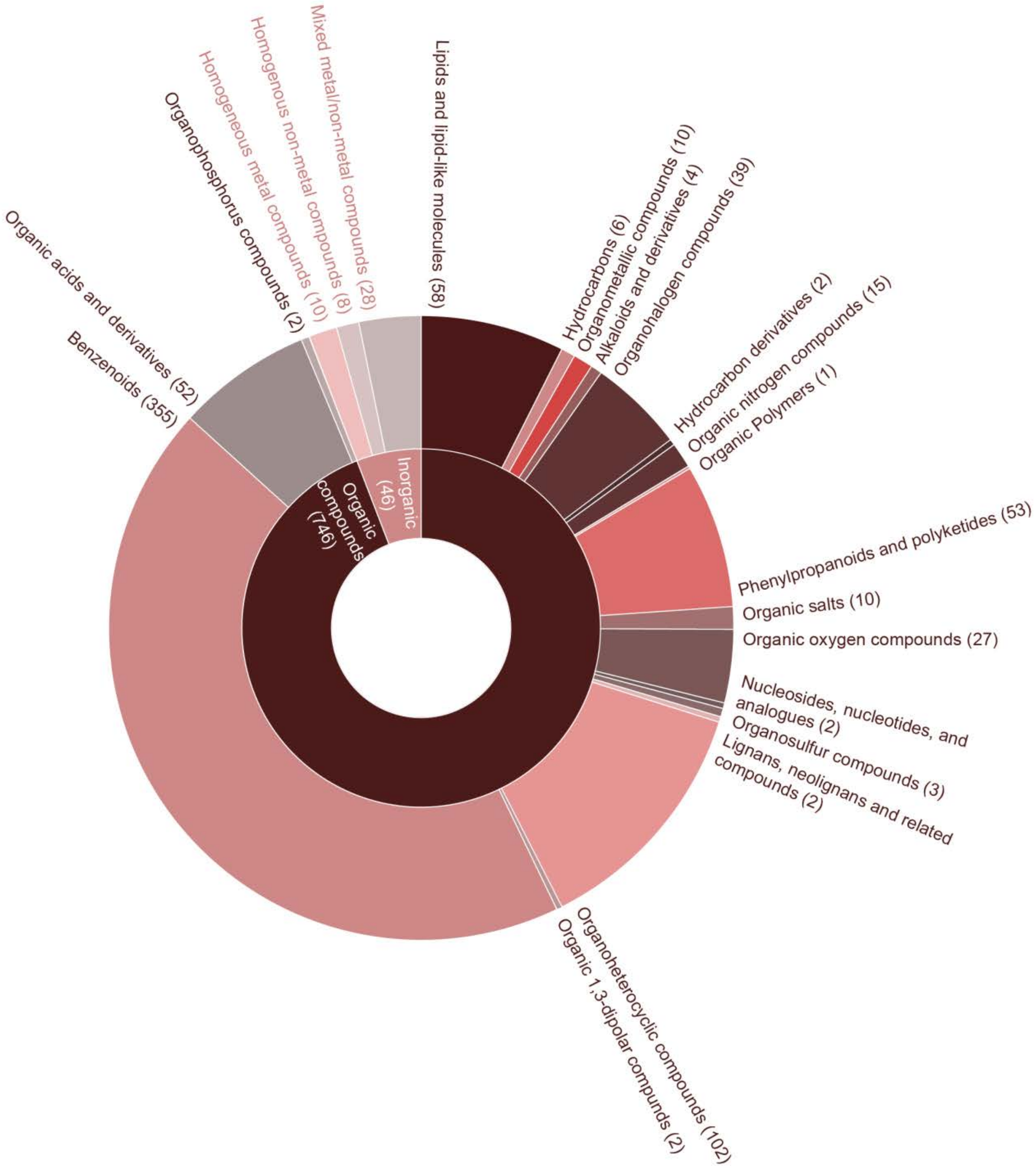


**Supplementary Figure S4:** Classification of the 792 EDCs in DEDuCT version 2.0 into chemical kingdoms and chemical super-classes using ClassyFire. Of the 792 EDCs, 746 are organic and 46 are inorganic compounds. The 746 organic EDCs can be further classified into 19 super-classes while the 46 inorganic EDCs fall into 3 super-classes. The number of EDCs in each super-class is reported within the parenthesis.

#### 1.2. Additional information on EDCs in DEDuCT version 2.0

In addition to experimental evidence, DEDuCT 2.0 also compiles diverse information for the 792 potential EDCs including two-dimensional (2D) and three-dimensional (3D) chemical structure, physicochemical properties, predicted ADMET properties, molecular descriptors, and experimentally inferred target genes from ToxCast database version August 2019 ("ToxCast Database (invitroDB)," 2018). We also provide a classification of the potential EDCs based on their environmental source into 7 broad categories and 48 sub-categories (Supplementary Figure S3). We also provide a hierarchical classification of the 792 potential EDCs based on their chemical structure information using ClassyFire (Djoumbou Feunang et al., 2016) (Supplementary Figure S4). Moreover, the final list of 792 potential EDCs were classified into 4 categories (I-IV) based on the type of supporting evidence for endocrine disruption in published experiments specific to humans or rodents (Supplementary Table S2). All the compiled information in DEDuCT can be downloaded in flat file format from the associated web server at: https://cb.imsc.res.in/deduct/download.

In sum, we provide an expanded list of potential EDCs in DEDuCT 2.0, which could assist academia, industry, and regulatory agencies to develop safer consumer products.

### 2. Description of chemical lists compiled from inventories, regulations or guidelines

#### 2.1. Substances in use (SIU) lists

##### 2.1.1. Plant protection products

We were able to obtain from public resources one SIU list in the category of ‘Plant protection products’.

- **List L1 - Active ingredients allowed in minimum risk pesticide products:** This list is published by the US Environmental Protection Agency (EPA). This list contains active plant protection ingredients that are considered minimum risk and are exempt from the requirements of the US Federal Insecticide, Fungicide, and Rodenticide Act. Plant protection products that contain these active ingredients, and meet the other criteria for minimum risk pesticides, do not need to be registered with the US EPA. This list is available for download from: https://www.epa.gov/minimum-risk-pesticides/active-ingredients-eligible-minimum-risk-pesticide-products

##### 2.1.2. Cosmetics and household products

We were able to obtain from public resources five SIU lists in the category of ‘Cosmetic and household products’.

- **List L2 - IFRA transparency list:** This list is published by International fragrance association (IFRA) which collectively represents the global fragrance industry. This list contains registered ingredients which are provided by fragrance companies. The list includes two types of ingredients, namely, fragrance ingredients and functional ingredients. Fragrance ingredients are basic chemical substances used for odor or malodor purposes while functional ingredients are chemical substances used to improve the performance and durability of fragrance. This list is available for downloaded from: https://ifrafragrance.org/initiatives/transparency/ifra-transparency-list
- **List L3 – EU list of colorants allowed in cosmetic products:** The cosmetic ingredient database (Cosing) is developed by the European Commission (EC) of the European Union (EU). The database contains several chemical lists based on different purpose of chemicals in cosmetics. One such list, the list of colorants allowed in cosmetic products, contains information on the colorants that are allowed in cosmetic products as per Annex IV of the Regulation (EC) No 1223/2009 of the European Parliament and its

council. This list is available for download from: https://data.europa.eu/euodp/en/data/dataset/cosing-list-of-colorants-allowed-in-cosmetic-products

- **List L4 – EU list of preservatives allowed in cosmetic products:** This is another list of chemicals in the Cosing database that are used in cosmetic products. This list contains information on preservatives allowed for use in cosmetic products as per Annex V of the Regulation (EC) No 1223/2009 of the European Parliament and its council. This list is available for download from: https://data.europa.eu/euodp/en/data/dataset/cosmetic-ingredient-database-list-of-preservatives-allowed-in-cosmetic-products
- **List L5 – EU list of UV filters allowed in cosmetic products:** This is another list of chemicals in the Cosing database that are used in cosmetic products. This list contains information on UV filters allowed in cosmetic products as per Annex VI of the Regulation (EC) No 1223/2009 of the European Parliament and its council. This list is available for download from: https://data.europa.eu/euodp/en/data/dataset/cosmetic-ingredient-database-list-of-uv-filters-allowed-in-cosmetic-products
- **List L6 - Consumer product ingredient database:** This dataset of chemicals in consumer products published by Goldsmith et al. (Goldsmith et al., 2014) contains information on unique chemical ingredients, consumer products and hierarchy of consumer product use categories. This dataset was created using material safety data sheet (MSDS) of products as provided by US retail corporation Walmart. The authors of the dataset had chosen the specific retail corporation based on its diverse product range and public access to its MSDS database. This list is available for download from: https://doi.org/10.1016/j.fct.2013.12.029

##### 2.1.3. Food additives and Food contact materials

We were able to obtain from public resources seven SIU lists in the category of ‘Food additives and Food contact materials’.

- **List L7 - Substances added to food (EAFUS):** This list is published by the US Food and Drug Administration (FDA) and was previously known as Everything Added to Foods in the United States (EAFUS). The list is an inventory of food additives, color additives and substances that are approved for specific use in food regulated by the US

FDA. This list is available for download from: https://www.accessdata.fda.gov/scripts/fdcc/?set=FoodSubstances

- **List L8 - FooDB:** FooDB is a manually curated open access database containing information on the chemical composition of commonly processed and unprocessed food. The database also contains information on food flavor, aroma molecules and food additives, which are further linked to their health effects. This list is available for download from: http://foodb.ca/
- **List L9 – JECFA food additives:** This list contains information on flavors, food additives, contaminants, toxicants and veterinary drugs that were evaluated by the Joint expert committee of the Food and Agriculture Organization (FAO) and World Health Organization (WHO) of the United Nations on food additives (JECFA). This expert committee meets twice a year to evaluate and summarize reports on food additives, contaminants and toxicants in the agricultural industry and veterinary drugs. This list is available for download from: http://apps.who.int/food-additives-contaminants-jecfa-database/search.aspx
- **List L10 – EU food flavorings database:** This EU database contains flavoring substances approved for use in food. This list was developed based on the Part I of Annex I of Regulation (EC) No 1334/2008, the EU regulatory document for food flavoring substances. This list is available for download from: https://webgate.ec.europa.eu/foods_system/main/?sector=FFL&auth=SANCAS
- **List L11 - EU plastic food packaging materials:** This EU list has relevance for the European Economic Area (EEA). This list is a commission regulation on the plastic materials that are used in food packaging. The materials are evaluated for their use in articles that are intended to come into contact with food, or already in contact with food, or reasonably expected to come into contact with food. This list is available for download from: https://eur-lex.europa.eu/eli/reg/2011/10/oj
- **List L12 - Pew list of food additives:** This list consists of direct and indirect food additives allowed in human food by US FDA with references to their toxicological information (Neltner et al., 2013). This database also discusses the limitations of the lists of food additives and available toxicity information. This list is available for download from: https://doi.org/10.1016/j.reprotox.2013.07.023

- **List L13 - ESCO list of non-plastic food contact materials:** This database of non-plastic food contact materials is published by the European Food Safety Authority Scientific Cooperation (ESCO) working group. This inventory consists of non-plastic chemicals found to be in contact with food such as coatings, colorants, cork and wood, paper and board, printing inks, rubber and silicones. This list is available for download from: https://doi.org/10.2903/sp.efsa.2011.EN-139

##### 2.1.4. Biocides

We were able to obtain from public resources one SIU list in the category of 'Biocides'.

- **List L14 – ECHA biocidal products:** This list is published by the European Chemicals Agency (ECHA) and gives a list of biocidal active substances and suppliers, according to Article 95 of the Biocidal Products Regulation (BPR), as amended by Regulation (EU) No 334/2014 of 11 March 2014. This list is available for download from: https://www.echa.europa.eu/regulations/biocidal-products-regulation/approval-of-active-substances/list-of-approved-active-substances

##### 2.1.5. Medicines and Medical devices

We were able to obtain from public resources one SIU list in the category of 'Medicines and Medical Devices'.

- **List L15 – US FDA inactive ingredient list:** This list published by US FDA contains information about inactive ingredients present in FDA-approved drug products. This database includes inactive ingredients in the final dosage forms of drug products. This list is available for download from: https://www.accessdata.fda.gov/scripts/cder/iig/index.cfm

##### 2.1.6. Miscellaneous lists

We were able to obtain from public resources two SIU lists in the 'Miscellaneous' category.

- **List L16 - Production of major chemicals year-wise in India:** This list is published by the Ministry of Chemicals and Fertilizers, Government of India and provides catalogs with data on the production of major chemicals in India. The data is presented biannually along with the production quantity of the chemicals. This list is available for download from: https://data.gov.in/catalogsv2?format=json&offset=0&limit=9&filters%5Bfield_ministr

y_department%3Aname%5D=Department+of+Chemicals+and+Petrochemicals&sort%5Bogpl_module_domain_name%5D=asc&sort%5Bcreated%5D=desc

- **List L17 – US EPA safer chemical ingredients list:** This list is published under the Safer Choice program of the US EPA. The chemicals in the list are arranged based on their functional-use class and categorized as 'low concern' based on the strength of the available supporting data. Products whose ingredients meet the Safer Choice criteria are allowed to carry the Safer Choice label. This list is available for download from: https://www.epa.gov/saferchoice/safer-ingredients#scil

#### 2.2. Substances of concern (SOC) lists

##### 2.2.1. Plant protection products

We were able to obtain from public resources two SOC lists in the category of 'Plant protection products'.

- **List L18 - List of banned pesticides in India:** This list is published by the Ministry of Agriculture and Farmers' Welfare, Government of India. This list is reviewed and updated to incorporate new findings regarding plant protection products. This list is available for download from: http://ppqs.gov.in/divisions/cib-rc/registered-products
- **List L19 - List of banned and restricted pesticide products in China:** This list is published by the Chemical Inspection and Regulation Service (CIRS). This is a list of plant protection chemicals that are restricted or prohibited in China. The list has been compiled from several documents regarding the prohibition or restriction of pesticides, released by several authorities, including the Ministry of Agriculture, People's Republic of China. This list is available for download from: http://www.cirs-reach.com/news-and-articles/List-of-Banned-and-Restricted-Pesticide-Products-in-China.html

##### 2.2.2. Cosmetics and household products

We were able to obtain from public resources one SOC list in the category of 'Cosmetic and household products'.

- **List L20 – EU list of substances prohibited in cosmetic products:** This EU list consists of chemical substances that are prohibited in cosmetic products as per Regulation (EC) No 1223/2009 of the European Parliament and of the Council of 30 November 2009 on cosmetic products. This regulation looks at cosmetic ingredients with human health as the focus. This list is available for download from: https://eur-lex.europa.eu/legal-content/EN/TXT/?uri=celex:02009R1223-20150416

##### 2.2.3. REACH chemicals

We were able to obtain from public resources two SOC lists categorized as 'REACH chemicals'. Here, REACH stands for Registration, Evaluation, Authorisation and Restriction of Chemicals.

- **List L21 – Restricted substances under REACH:** This list published by the ECHA consists of substances (on their own, in a mixture or in an article) that are restricted or banned for manufacturing or marketing in the EU. This list is the Annex XVII of the EU chemical regulation known as REACH. This list is available for download from: https://echa.europa.eu/substances-restricted-under-reach
- **List L22 - SVHC under REACH:** This list is published by the ECHA. The substance of very high concern (SVHC), also called the Candidate list, is a list of hazardous substances which will slowly be replaced by safer alternatives in the EU. Chemicals are reviewed for addition to this list when ECHA or any EU member state finds a substance to be SVHC. This list is available for download from: https://echa.europa.eu/substances-of-very-high-concern-identification-explained

##### 2.2.4. Environment and Water Quality

We were able to obtain from public resources five SOC lists in the category of 'Environment and Water Quality'.

- **List L23 - NPI Australia:** This list is published by National Pollutant Inventory (NPI) Australia. Based on potential impact of chemical substances on health and environment, the NPI lists 93 chemicals as priority substances along with their threshold. This list is available for download from: http://www.npi.gov.au/substances/substance-list-and-thresholds
- **List L24 – Singapore list of controlled hazardous substances:** This list captures Schedule 2 of the Environmental Protection and Management Act (EPMA) of Singapore which lists hazardous substances that are regulated in manufacturing and import. This list is available for download from: https://www.nea.gov.sg/docs/default-source/default-document-library/hs--table-1.pdf

- **List L25 - Ozone-depleting substances in India:** This list is published by the Ministry of Environment and Forests, Government of India. Based on sections 6, 8 and 25 of the Environment (Protection) Act 1986, the rules for regulating ozone-depleting substances were published, and in Schedule-I, the list of ozone-depleting substances has been notified. This list is available for download from: https://npcb.nagaland.gov.in/wp-content/uploads/2016/03/Ozone-Rules-2000.pdf
- **List L26 - EWG tap water database:** This database by the non-profit organization Environment Working Group (EWG) contains drinking water quality data for nearly 50000 community water systems in USA. EWG has curated this database using water testing records maintained by the government water authorities, and US EPA's test data on unregulated contaminants. This list is available for download from: https://www.ewg.org/tapwater/
- **List L27 - Human Indoor Exposome database:** This database is a compilation of 511 compounds found in indoor dust which were compiled using data mining (Dong et al., 2019). This list was further analyzed using the ToxCast database of US EPA. The list is available for download from: https://doi.org/10.1021/acs.est.9b00280

##### 2.2.5. Workers' regulations

We were able to obtain from public resources one SOC list in the category of 'Workers' Regulations'.

- **List L28 – US OSHA list:** This list published by the US Department of Labor Occupation Safety and Health Administration (OSHA) lists toxic and reactive highly hazardous chemicals that are of concern under the Occupational Safety and Health Standards. This list provides the chemical name, the CAS identifier and the threshold quantity of each chemical. This list is available for download from: https://www.osha.gov/laws-regs/regulations/standardnumber/1910/1910.119AppA

##### 2.2.6. Miscellaneous

We were able to obtain from public resources eight SOC lists in the 'Miscellaneous' category.

- **List L29 - SIN List:** This list is published by the non-profit organization ChemSec. The Substitute It Now! (SIN) list is a list of hazardous chemicals that are widely used across the globe and pose threat to human health and environment. This list is developed based on the EU REACH criteria for SVHC. The SIN list also provides information on available alternative chemicals to substitute hazardous chemicals. This list is available for download from: https://sinlist.chemsec.org/

- **List L30 - Toxic chemicals restricted to be imported or exported in China:** This list contains hazardous chemicals severely restricted to be imported into or exported from China under The Chinese Ministry of Environmental Protection order No. 22 and updated in 2014. This list is available for download from: http://www.cirs-reach.com/China_Chemical_Regulation/Registration_of_import_export_of_toxic_chemicals_in_China.html
- **List L31 - IARC monographs on carcinogens:** This list is published by the International Agency for Research on Cancer (IARC), a part of the WHO of the United Nations. Since 1970, IARC has published scientific reviews on the carcinogenicity of various substances that humans are exposed to in a variety of ways. The chemical agents reviewed in the monographs have been classified into four groups: (a) Group 1 - Carcinogenic to humans, (b) Group 2A - Probably carcinogenic to humans, (c) Group 2B - Possibly carcinogenic to humans, and (d) Group 3 - Not classifiable as to its carcinogenicity to humans. This list is available for download from: https://monographs.iarc.fr/agents-classified-by-the-iarc/
- **List L32 - Schedule 1 hazardous chemical list in India:** This list is published by the Ministry of Environment and Forests, Government of India. Under the notification on 'Manufacture, Storage And Import Of Hazardous Chemical Rules, 1989', Part II of Schedule 1, titled 'Indicative Criteria and List of Chemicals', lists the hazardous and toxic chemicals that fall under this regulation. This list is available for download from: http://moef.gov.in/wp-content/uploads/2019/08/SCHEDULE-I.html
- **List L33 - Schedule 3 hazardous chemical list in India:** This list is published by the Ministry of Environment and Forests, Government of India. Under the notification on 'Manufacture, Storage And Import Of Hazardous Chemical Rules, 1989', and as per Schedule 3 in Part I, the 'List of Hazardous Chemicals for Application of Rules 5 and 7-15' are mentioned. Rule 5 pertains to the reporting of major accidents. Rules 7-15 pertain to notification of the sites of use of the chemicals in Schedule 3, submission and update of safety reports, and management plans in case of major accidents. This list is

available for download from: http://moef.gov.in/wp-content/uploads/2019/08/SCHEDULE-3.html

- **List L34 – NZ EPA priority chemical list:** This list is published by the Environmental Protection Authority (EPA) of New Zealand (NZ). This is a list of chemicals of potential concern according to either NZ EPA or regulatory bodies in other parts of the world. The chemicals are screened using Flexible Reassessment Categorisation Screening Tool (FRCaST). This list is available for download from: https://www.epa.govt.nz/industry-areas/hazardous-substances/chemical-reassessment-programme/priority-chemicals-list/
- **List L35 – ECHA list of chemicals in Annex I:** This list published by ECHA consists of individual chemicals or groups of chemicals that come under the Prior Informed Consent (PIC) Regulation, which regulates the export and import of hazardous substances. This list is divided into three parts based on the regulatory subcategory the chemical fall under. This list is available for download from: https://echa.europa.eu/information-on-chemicals/pic/chemicals
- **List L36 - PACSs list Japan:** This list is published by the National Institute of Technology and Evaluation (NITE) Japan. Priority Assessment Chemical Substances (PACSs) database was developed based on evaluations relevant to human health effects and/or ecological effects. The database gives the list of evidences, chemical identifiers and classification of chemical substances. This list is available for download from: https://www.nite.go.jp/en/chem/chrip/chrip_search/intSrhSpcLst?slIdxNm=&slScNm=RJ_01_020&slScCtNm=&slScRgNm=&ltCatFl=&slMdDplt=2&ltPgCt=100&stMd=
